# Design, Expression, and Purification of a Soluble Form of the Retina-Specific Membrane Transporter, ABCA4

**DOI:** 10.1101/2025.05.22.655561

**Authors:** Albtool A. Alturkestani, Jazzlyn S. Jones, Senem Cevik, Esther E. Biswas-Fiss, Subhasis B. Biswas

**Affiliations:** Department of Medical and Molecular Sciences, University of Delaware, College of Health Sciences, Newark, DE 19716; Ammon Pinizzotto Biopharmaceutical Innovation Center, 590 Avenue 1743 • Newark, DE 19713

**Keywords:** Membrane transporter, Visual diseases, Protein engineering, Water-soluble transporter protein, ABCA4 protein

## Abstract

The ATP-binding cassette transporter A-subfamily member, ABCA4, is highly expressed in rod and cone photoreceptors in the retina, where it transports *cis-* and *trans-*retinal and is indispensable for vision. Genetic mutations in the ABCA4 gene lead to a wide range of inherited retinal degenerative diseases, including Stargardt disease (STGD1) and autosomal recessive cone-rod dystrophy. It is an integral membrane protein with twelve transmembrane α-helices that complicates studies with the full-length ABCA4 transporter. We have engineered the full-length ABCA4 by transforming its membrane helices, creating a soluble homolog (ABCA4s). Most hydrophobic residues in the membrane helices were substituted with structurally compatible but hydrophilic residues. The re-engineered ABCA4s was expressed in insect cells, and it was found in the cytosolic extract, which was purified by immunoaffinity chromatography. Purified ABCA4s was enzymatically active, all-trans-retinal stimulated its ATPase activity, and its activity remained stable.

**Significance:** ABCA4 is a 12-pass transmembrane protein that plays essential roles in the human retina and multiple visual diseases. Historically, the purification of ABCA4 and other large membrane proteins has relied on detergent-based purification, which renders the protein’s enzymatic activity highly unstable. We describe here the design of a truly soluble analog of ABCA4 with stable enzymatic activities. Its 12 transmembrane helices were transformed using selective amino acid substitution, and deleterious substitutions were carefully avoided. This soluble form will pave the way for mechanistic studies of the enzyme and its disease-causing genetic variants. The methodology described here should be widely applicable to other complex membrane proteins facilitating their studies without the need for reconstitution in lipids.

## Introduction

Membrane transporter proteins play pivotal roles in cellular metabolism by transporting various molecules across the cell and subcellular membranes (1, 2). One such transporter protein family is the ATP-binding cassette (ABC) transporters (1, 3, 4). Forty-nine ABC genes have been identified to date and are categorized into eight subfamily groups, A to H, with very diverse functional roles (4–6). The ABCA transporter subfamily has 13 members expressed in different cell tissue types. Mutations and dysfunctions of these transporter proteins are involved in various diseases (4). The fourth member of the ABCA subfamily is the ABCA4 transporter protein, also known as ABCR or Rim protein. ABCA4 is localized in the rod and cone outer segment disc membrane of photoreceptor cells *(SI Appendix, Fig. S1)*. It functions as a transporter for *cis* and *all-trans*-retinal (ATR) and, thus, is essential for vision. In the visual cycle of the retina, *cis*-retinal bound to rhodopsin isomerizes to *trans* by absorbing a photon. ABCA4 translocates ATR to the lumen. After its isomerization to *cis*, *11-cis* retinal is imported to the cytoplasmic region again by ABCA4 (7, 8). Purified ABCA4 from bovine rod outer segments identified *11-cis* and *all-trans*-retinal as ligands to stimulate ATPase activity (9).

ABCA4 shares structural topology with all members of the ABCA subfamily that include two extracellular domains (ECDs), two transmembrane domains (TMDs) that have 12 α-helix segments located within the lipid bilayer, and two nucleotide nucleotide-binding domains (8). Genetic variants of ABCA4 may lead to dysfunction in the transport of *cis* and *trans*-retinal across disc membranes leading to various forms of visual diseases (4, 10). At present, more than 2761 unique ABCA4 variants have been reported. Missense, frameshift, stop, and splice site variants in ABCA4 are linked to inherited diseases such as Stargardt disease (STGD1), cone-rod dystrophy, fundus flavimaculatus, and autosomal retinitis pigmentosa type 19 (4, 11, 12). Molecular studies showed that variants in the ABCA4 gene can disrupt the protein’s function (13–16). Understanding the structure and function of ABCA4 protein is essential to delineate how it transports substrates across the membrane and how missense mutations in ABCA4 lead to inherited retinal disease. Assessing defects in ABCA4 mutants *in vitro* has been challenging due to its large size and membrane association. In general, ABCA proteins are very large in size and are difficult to express in recombinant form. Extraction and purification require harsh detergents that disrupt the membrane lipid bilayer to effect solubilization, and this process adversely affects protein stability (17–19). Such problems are associated with most membrane proteins and are not unique to ABCA4 or the ABCA family of transporters (20, 21).

DeGrado et al. solubilized Phospholamban by replacing certain hydrophobic amino acid residues in the membrane-embedded α-helix of the protein (22). Thereafter, several groups attempted to create soluble analogs of integral membrane proteins using computational tools to aid in the design. A common theme of these studies has been the substitution of hydrophobic residues of transmembrane helices with suitable hydrophilic helix-forming amino acids, for example, leucine (Leu, L) to glutamine (Gln, Q), Glutamic acid (Glu, E), or Lysine (Lys, K). Several transmembrane proteins were solubilized using these approaches, such as the potassium ion-gated channel KcsA, nicotinic acetylcholine receptor (AChR), mu opioid receptor, and G-protein coupled receptors (22–26). In 2018, Zhang *et al.* proposed the QTY substitution approach to convert leucine (Leu, L) to glutamine (Gln, Q) or asparagine (Asn, N), valine (Val, V) and isoleucine (Ile, I) to threonine (Thr, T), and phenylalanine (Phe, F) to tyrosine (Tyr, Y) in the membrane α-helices (27). The substitution of a hydrophobic amino acid for a hydrophilic amino acid is based on the comparable electron density map and a neutral side chain of each amino acid. Thus far, the application of the amino acid substitution approach has been successful in smaller transmembrane proteins. We explored the possibility of applying a similar approach to a protein as large as ABCA4 with 2,273 amino acids and 12 transmembrane domains. An initial attempt to apply the QTY approach to ABCA4 did not work. Thereafter, we developed an approach of limited substitution of the membrane hydrophobic residues based on structural analysis and databases of genetic variant pathogenicity. The modified membrane-free ABCA4 (ABCA4s) is soluble and does not require refolding. This strategy may be helpful to solubilize other large transmembrane proteins of biomedical significance.

## Results

ABCA4 is one of many large membrane proteins that are physiologically essential and clinically important. However, these proteins are generally challenging to study because of the large size and membrane association. In addition, membrane proteins are often difficult to purify and frequently unstable in purified form (17–21). Consequently, there have been efforts to create soluble analogs of membrane proteins to make them amenable to biochemical studies (22–27). We have used this concept to create a soluble form of ABCA4 protein.

### Application of the QTY approach to ABCA4s

We first attempted to utilize the recently published QTY methodology to convert ABCA4 into a soluble form (27). This approach directed us to create a synthetic codon-optimized ABCA4_QTY_ DNA sequence with mutations in all 12 TMDs and the two smaller membrane helices EH3-EH4 *(SI Appendix, Fig. S2 and Table 1)*. We replaced 133 hydrophobic residues (Leu, Val, Ile, and Phe) in the 12 transmembrane α-helices and EH3-EH4 that were changed as follows: Leu → Gln, Val/Ile → Thr, and Phe → Tyr. We created the ABCA4_QTY_ DNA sequence with these modifications for expression in a baculovirus system. A synthetic baculovirus transfer vector with an expression cassette containing the ABCA4_QTY_ ORF was synthesized, and the virus was generated following the standard protocol of Multibac^TM^ system (Geneva-Biotec, Geneva, Switzerland) (28–30). The virus was amplified in insect cells (V1 virus). The amplified virus was used to infect High Five^TM^ (Hi5) cells for expression of the protein. We made several attempts with this construct but could not express the ABCA4_QTY_ protein. We could not determine the reasons for the lack of expression with this construct. Therefore, we explored possible modifications of the procedure to overcome the lack of expression.

**Table 1.** Summary of amino acid substitutions in the 12 transmembrane α-Helical segments of ABCA4s. A list of substitutions in each helix is presented here. Pathogenic amino acid substitution(s) that were omitted are listed as “No substitution” and the sources of these variants are listed in the last column.

| TMS | L > Q | L > N | I > T | V > T | P > Y | Start and End of each TMS (amino acid number) | Total QNTY/TMS full-length | No Substitutions | Pathogenic Residues ClinVar/Literature |
| --- | --- | --- | --- | --- | --- | --- | --- | --- | --- |
| 1 | 2 | 2 | 2 | 4 | 2 | 23-42 | 12/20 | 2 | ClinVar: L29R, S34_L35del |
| 2 | 0 | 1 | 3 | 4 | 1 | 657-676 | 9/20 | 1 | Garces *et al* (2018): L661R |
| 3 | 1 | 2 | 4 | 0 | 4 | 696-718 | 11/23 | 0 | - |
| 4 | 3 | 2 | 1 | 0 | 6 | 733-755 | 12/23 | 2 | ClinVar: L750P<br>Garces *et al* (2018): L737 |
| 5 | 1 | 2 | 2 | 0 | 2 | 762-782 | 7/21 | 1 | ClinVar/Garces *et al* (2020): V767D |
| 6 | 3 | 3 | 0 | 0 | 0 | 837-853 | 6/17 | 1 | Garces *et al* (2018): V849A |
| 7 | 1 | 3 | 2 | 3 | 2 | 1,376-1,393 | 11/18 | 0 | - |
| 8 | 1 | 1 | 3 | 5 | 3 | 1,681-1,702 | 13/22 | 0 | - |
| 9 | 0 | 2 | 3 | 3 | 2 | 1,727-1,747 | 10/21 | 0 | - |
| 10 | 2 | 2 | 1 | 2 | 0 | 1,762-1,782 | 7/21 | 1 | ClinVar: L1763P |
| 11 | 0 | 2 | 3 | 1 | 1 | 1,789-1,810 | 7/21 | 0 | - |
| 12 | 2 | 3 | 0 | 5 | 2 | 1,876-1,896 | 12/21 | 0 | - |

### Development of an alternative approach

In the second attempt, the cryoEM structure of ABCA1, another member of ABCA transporter family, became available and we first aligned the ABCA4 sequence to the ABCA1 cryo-EM structure (31). Based on this alignment, we could streamline the number of residues needed to be substituted from 133 to 125 to solubilize the ABCA4 protein (ABCA4s). The substitutions in the final TMDs with sequences are shown as a synopsis in Figure 2, table 1. However, the overwhelming numbers of substitutions are still a concern as any of these substitutions could adversely affect the structure of such a large protein. Therefore we explored avenues to identify and eliminate possible structurally damaging substitutions. First, we analyzed the mutations using PolyPhen-2 pathogenicity prediction tool and PyMol softwares (32, 33). Second, we cross-checked our tentative substitution list against ABCA4 clinical databases (ClinVar and LOVD) to identify substitutions that could lead to structural and/or functional defects (34–36). Third, we also considered pathological variants of ABCA4 reported in the literature but not yet included in various ABCA4 variant databases.

### Bioinformatics prediction of pathogenic residues

We have analyzed all 125 substitutions by PolyPhen-2 and PyMol software for possible structural damages associated with the substitution (32, 33). Several of the QTY mutations turned out to be damaging. In the mutations with L→ Q substitution, we examined whether L → N substitution was more suitable. As the Leu electron density map is similar to Gln as well as Asn (Figure 1 A), L→N appeared to be a suitable substitution of L→Q. Thus, we considered Leu → Asn substitution as an alternative in certain cases for L→ Q substitution (*SI Appendix, Table 2)*. We changed seven L→ Q substitutions to L→ N substitutions based on the PolyPhen-2 analysis. In addition, we changed 18 L→ Q substitutions to L→ N substitutions based on the PyMol analysis.

**Fig. 1.**
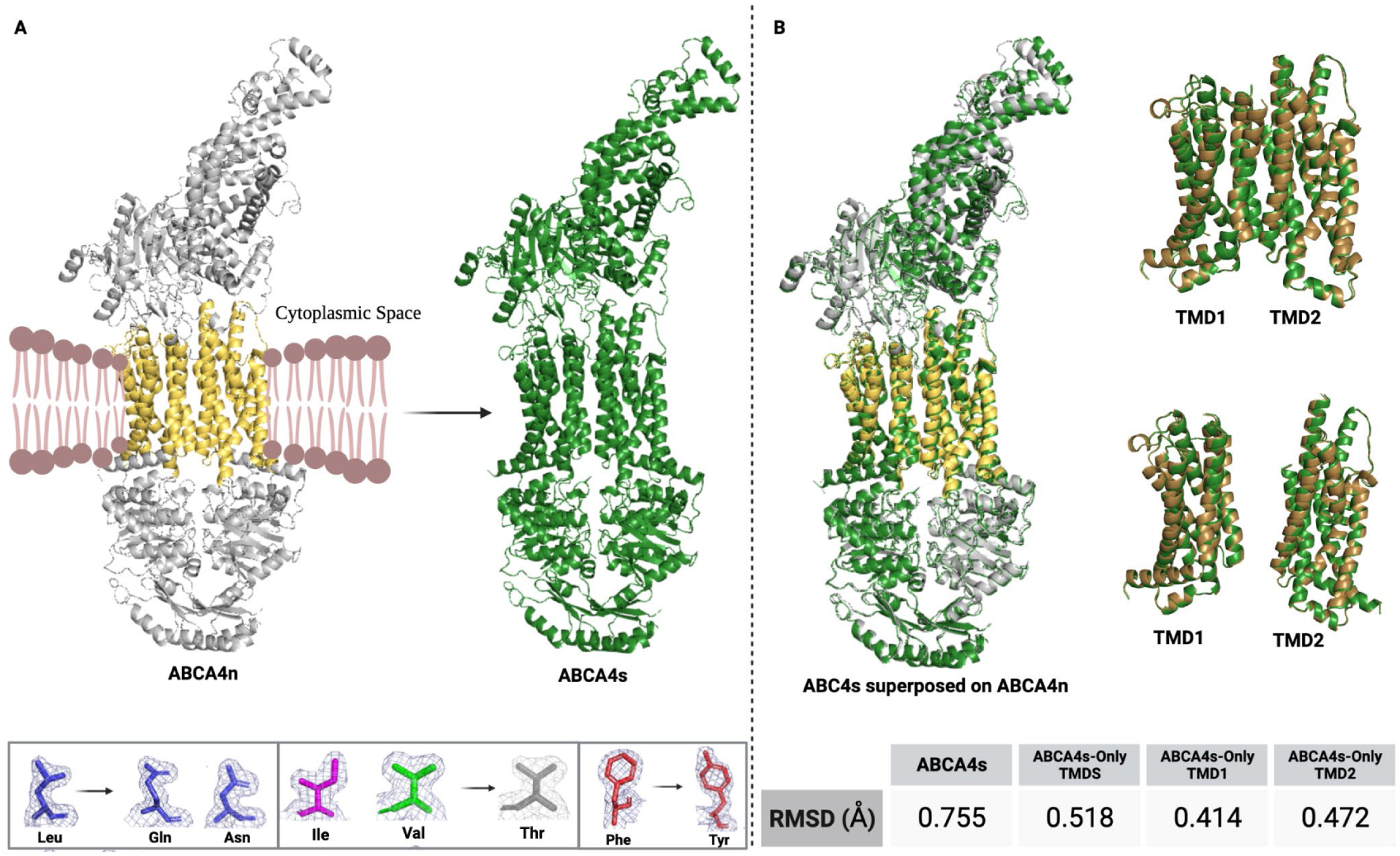
Alphafold-2 analysis of native ABCA4n and soluble ABCA4s. **A.** Conversion of the native ABCA4n protein to its soluble form ABCA4s. Alphafold-2 models of ABCA4n and ABCA4s are shown. Electron density maps of L, I, V, F, Q, N, T, and Y are shown that form the basis of the amino acid sustitution. **B.** Superimposition of the ABCA4n and ABCA4s models: full-length proteins (left), transmembrane domains (TMDs) (top-right), and separately modeled TMD1 and TMD2 (bottom-right). Gray and yellow represent the ABCA4n, and green represents the ABCA4s structures. The models were visualized and analyzed in PyMOL2. (Created with BioRender.com).

### Clinically established pathogenic mutations in the ABCA4 TMDs

ABCA4 is associated with several inherited retinal degenerative diseases and, thus, has been a subject of various genetic and clinical investigations over decades (4, 11–14). To date, >2700 genetic variants have been identified in the ABCA4 gene and are described in mutational databases. Many of these variants are listed in the clinical genetic database, ClinVar (34, 37). QTY substitutions where residues were identical to ABCA4 reported pathogenic variants were deemed unsuitable for substitution and avoided (34, 37). As a result, four intended QTY mutations were eliminated, as shown in Table 1.

### Mutations in the ABCA4 TMDs in the published literature

Next, we considered four of the 125 QTY mutations, which were identified as structurally deleterious in published clinical reports (14, 38). These are L661, L737, V767, and V849 (Table 1, row 10). As a result, substitutions involving these residues were not considered.

Overall, out of 125 intended QTY mutations, we carried out 117 QNTY substitutions as follows 16 L→ Q, 25 L→ N, 24 I → T, 27 T→ Q, and 25 F→ Y (Figure 2). Finally, we used the TMHMM-2.0 server to evaluate the final design of the ABCA4s to ascertain hydrophobic transmembrane helices are removed *(SI Appendix, Fig. S3)*. The designed ORF was cloned into a baculovirus transfer vector under a Polyhedrin baculovirus promoter.

**Fig. 2.**
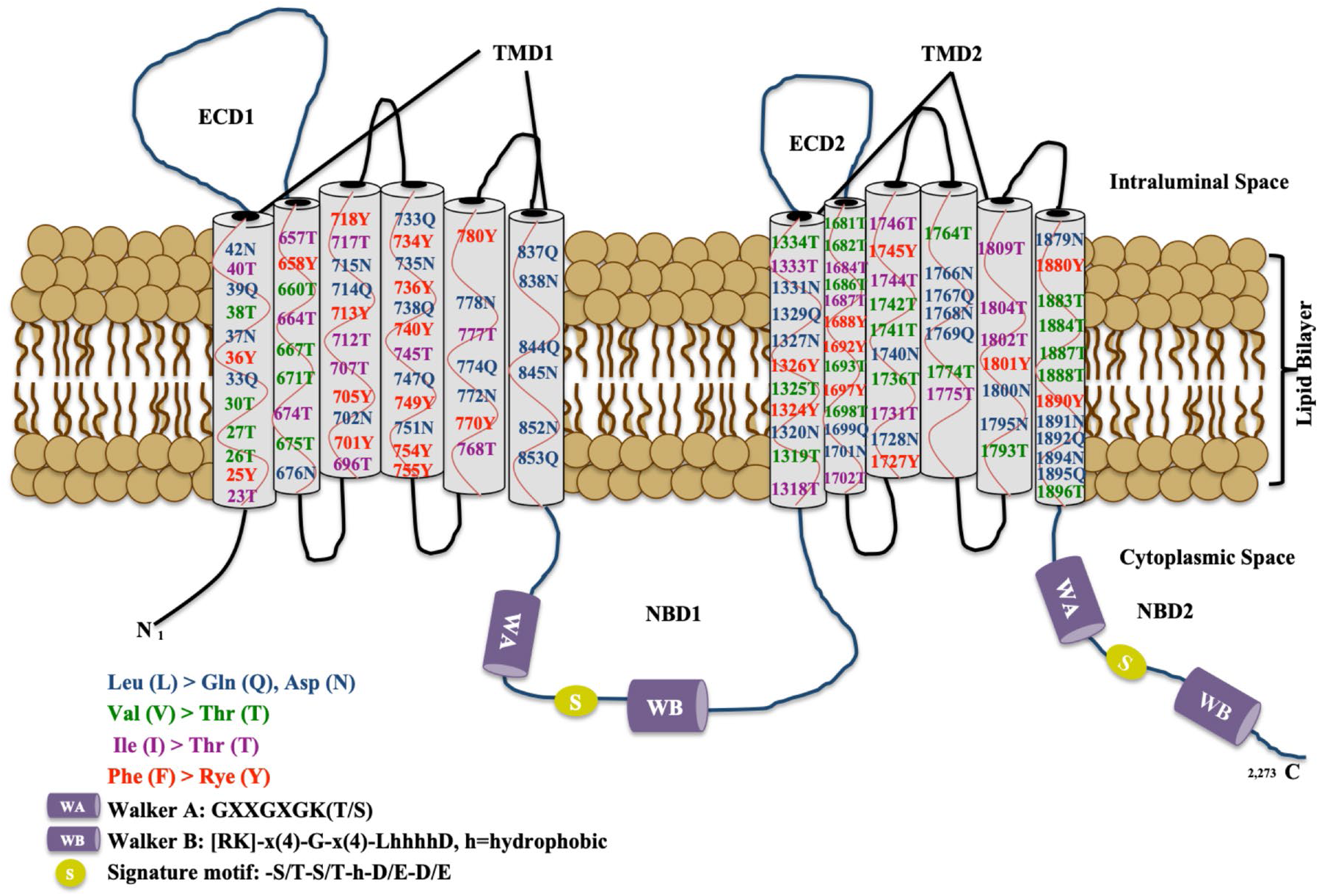
Amino acid substitution map of ABCA4s. ABCA4 protein in membrane-bound form with two extracellular domains (ECDs), two transmembrane domains (TMDs) each containing six α-helices with indicated substitutions (Color code: L → Q/N is shown in blue, I → T is presented in pink, V → T is colored in green, and F → T is shown in red). Numbers indicate the sequence location of QNTY substitutions.

### *In silico* modeling and protein structure analysis

We generated computational models of the native (ABCA4n) and soluble (ABCA4s) proteins in AlphaFold2 as full-length, only for the TMDs, and only TMD1 and TMD2 separately (39, 40). All models were generated in high confidence *(SI Appendix, Fig. S4)*. ABCA4s’ protein structure retained all the α-helices constituting the transmembrane domains. We compared the predicted ABCA4n and ABCA4s structures by superimposition. We found a high degree of structural similarity and low RMSD between the native and the soluble *in silico* models for all three modeling approaches (Figure 1B). The RMSD score was 0.944 Å for the full-length ABCA4s alignment with the native ABCA4 model, and 0.518 Å for the only-TMD models. Although both soluble protein structures exhibit remarkable structural alignment with the native models, the TMD2 alignment score was slightly higher in the individual TMD modeling; RMSD_TMD1_: 0.414 Å, RMSD_TMD2_: 0.472 Å.

We found no significant difference in the molecular weight and isoelectric focusing point (pI) between the two proteins. The molecular weight of the ABCA4n is 255943.83, and the pI is 6.2, whereas ABCA4s mol. wt. is 256370.92 and the pI is 6.2.

### Expression, and purification of ABCA4 soluble form (ABCA4s)

The synthetic ABCA4s ORF was cloned into pBV plasmid under the control of polyhedrin promoter. ABCA4s protein was expressed in Hi5 cells (from *Trichoplusia ni),* and expression of ABCA4s was confirmed by SDS-PAGE and Western blot analysis (Figure 3A). As shown in figure 3A, ABCA4s produced in high yield at the expected size above 250kDa.

**Fig. 3.**
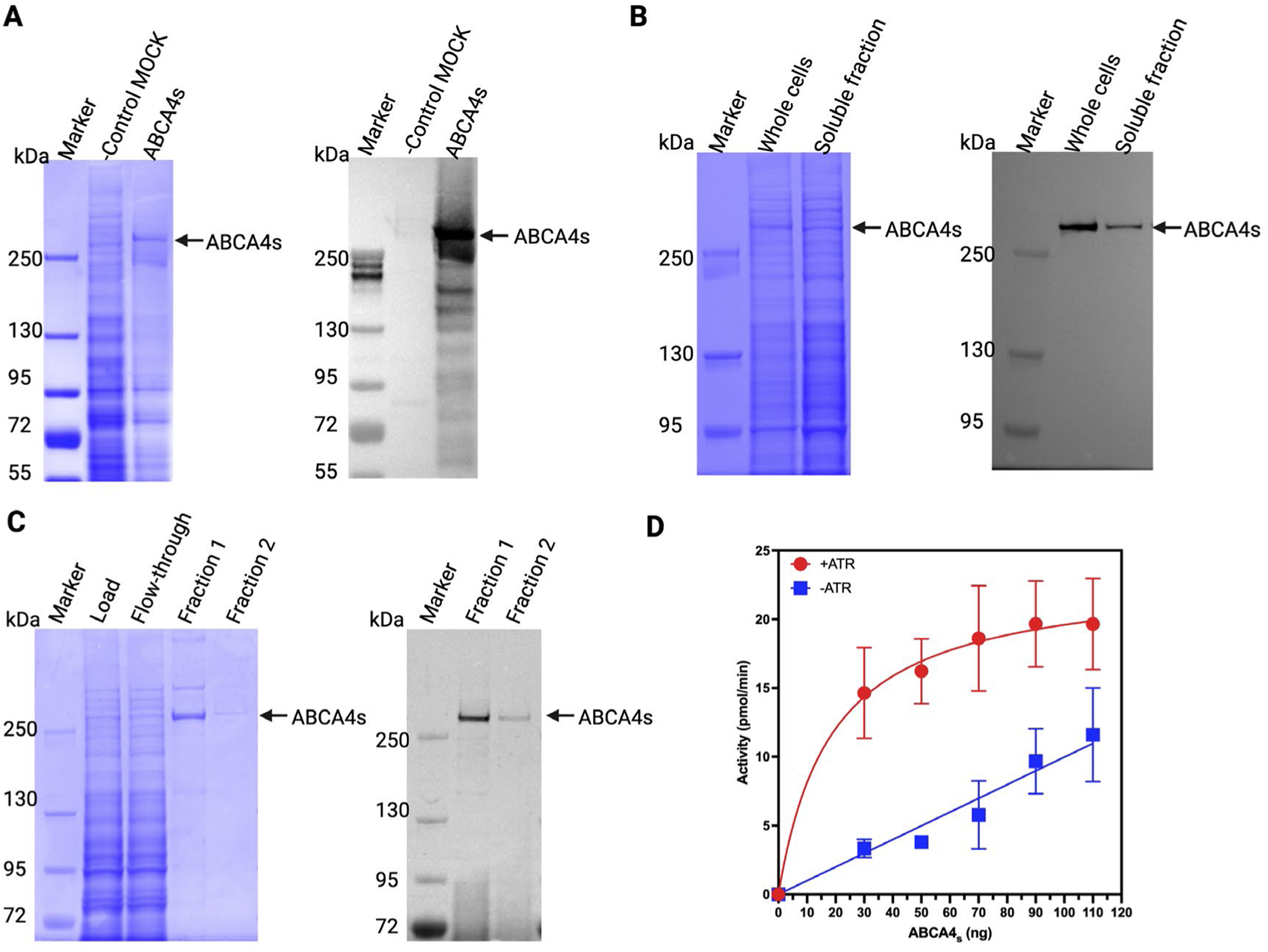
Analysis of solubility, purification, and biological activities of ABCA4s. **A.** SDS-PAGE and Western blot analysis of 62 hours post-infection Hi5 cells infected with ABCA4s expression virus or a blank virus (MOCK). In A-C, blue pictures are of Coomassie-stained gels, whereas black and white illustrations are of Western blots. **B.** SDS-PAGE and Western blot analysis of cytoplasmic extract of Hi5 cells as indicated. **C.** SDS-PAGE and Western blot analysis of purified ABCA4s protein by immunoaffinity chromatography. **D.** ATPase activity of ABCA4s was measured in the presence and absence of ATR using a luminescence-based ATPase assay. The error bars represent the standard deviations of triplicate runs.

The soluble proteins from infected Hi5 cells were extracted using an insect cell extraction reagent, IPER buffer (Thermo Fisher Scientific, Waltham, MA). The whole cells were analyzed along with the soluble protein extract. Upon SDS-PAGE and Western blot analyses, we observed ABCA4s in the cytoplasmic extract (Figure 3B). The soluble ABCA4s was then purified by immunoaffinity chromatography using anti-ABCA4 antibody and magnetic Protein A/G beads. ABCA4s in the eluted fractions were analyzed using SDS-PAGE. ABCA4s appeared to be of >90% purity in SDS-PAGE (Figure 3C). Next, we analyzed the ABCA4s fractions using western blot analysis and an antibody against ABCA4. As expected, a ∼250 kDa protein band was visible in the Western blot that was also visible in the Coomassie-stained SDS-PAGE gel. However, we observed a minor higher molecular weight band on the Coomassie-stained gel that was not detected in the Western blot, which indicated that the higher protein band was likely an impurity from insect host cells (Figure 3C).

### Functional analysis of ABCA4s

ABCA4 is an enzyme with specific ATPase activity, stimulated by *all-trans-*retinal (ATR) (9). We measured the ATP hydrolysis in the presence and absence of ATR to test the protein’s functionality. ATPase assays were carried out as described in the presence of 0.5 mM ATP and 40 μM ATR. ATR stimulated the activity approximately two-fold, comparable to the native ABCA4 in previous studies (9, 17, 43). In summary, purified ABCA4s enzyme was enzymatically active, stimulated >two-fold by the ATR and the activity remained stable at 4°C for at least 72 hours, and three-months at -80°C (Figure 3D). In the presence of ATR, the specific ATPase activity was ∼420 nmol/min/mg.

## Discussion

ABCA4 is a retina-specific transporter protein that transports *cis-* and *all-trans*-retinal in the human retina and is essential for vision. With more than 2,700 identified clinical variants, *ABCA4* pathogenic mutations are a leading cause of autosomal recessive inherited retinal disorders, including Stargardt macular degeneration, cone-rod dystrophy, and retinitis pigmentosa. Because of its large size, this 12-pass integral membrane protein has been difficult to study *in vitro* (17–19). Therefore, we explored the possibility of creating a full-length soluble analog of the ABCA4 protein using the recently published QTY approach (27). Initially, we identified 133 hydrophobic Leu, Ile, Val, and Phe residues in the 12 trans-membrane helices that needed to be substituted by Gln, Thr, and Tyr, respectively. However, the protein expression was undetectable when this analog if ABCA4 was synthesized and the synthetic ABCA4qty gene was expressed in insect cells using the baculovirus expression system. We hypothesized that one or more of these substitutions might be structurally deleterious.

We developed an algorithm to optimize the design of the soluble form of ABCA4 utilizing the QTY methodology. This approach involved careful assessments of each nucleotide/amino acid substitution using available structural tools and genetic databases to identify possible deleterious mutations that may cause dysfunction and loss of solubility. Ultimately, this reduced the number of substitutions from 133 to 117, creating the engineered ABCA4 protein, ABCA4s. This methodology also conferred a downstream experimental advantage because ABCA4s does not harbor any mutation in sequence other than the membrane helices, thereby retaining wild-type functional domains enabling it to be used as a platform in future studies to assess disease-associated *ABCA4* variants. ABCA4s could thus help close a major gap in visual disease research because the enzymatic effects of hundreds of ABCA4 disease-associated variants currently remain unknown.

As a clinically significant protein, many of ABCA4’s well over 2700 genetic variants have been correlated with protein malfunctions leading to various visual disorders (11, 12, 34, 37, 44). In our study, each of the 125 intended substitutions in the ABCA4 protein were analyzed by (i) PolyPhen-2 and PyMOL software which predicts the impact of an amino acid substitution on the structure of a protein (32, 33), (ii) reports in that are published in the NCBI ClinVar database (34, 37), and (iii) ABCA4 pathogenic mutations that are published in the literature (14, 38). In this approach, we found 33 of the 125 substitutions were problematic. Interestingly, 25 of these substitutions were Leu → Gln. Therefore, we elected to switch these substitutions to Leu → Asn. For the other eight substitutions, we chose to omit them altogether. In summary, we selected 117 substitutions (16 Leu → Gln, 25 Leu → Asn, 24 Ile → Thr, 27 Val → Thr, and 25 Phe → Tyr) (Table 1, Figure 2) in the final design of the ABCA4s protein.

Recently, cryo-EM structures of ABCA4 in the presence of the N-Ret-PE ligand (NRPE) became available which showed the residues that interacted with NRPE (45). The ATR moiety of NRPE is located between four aromatic residues (W339, Y340, Y345, and F348) in an extended loop of ECD1, which were not substituted (45). The β-ionone ring of the ATR moiety is integrated between TMD8 and TMD11 within five hydrophobic residues (I1812, L1815, L1674, S1677, and Y345) and further stabilized through the interactions with the aromatic residues (W339, Y340, and F348), and interestingly none of these residues were substituted in our design of ABCA4s (41).

Although previously reported QTY protein modifications have used *E. coli* expression systems, ABCA4 is a very complex human protein and thus, we chose to express ABCA4s in insect cells. We considered the possibility that post-translational modifications in insect cells may also help with solubility and biological functions. The ABCA4s expressed at high level in insect (Hi5) cells (Figure 3A). Upon extraction, ABCA4s was primarily found in the cytoplasmic fraction without any detergent indicating its solubility (Figures 3A, 3B). ABCA4s was purified to near homogeneity by immunoaffinity chromatography using an immobilized antibody against epitopes of ABCA4 (Figure 3C).

Purified ABCA4 protein is characterized *in vitro* by two distinct activities (9). First, it is an ATPase. Second, it binds to *all-trans*-retinal and the binding is detected by its stimulation of the ATPase activity. We examined the ATPase activity of ABCA4s and the stimulation by ATR. The ATPase activity of ABCA4s in the presence and absence of ATR is shown in Figure 3D. ATPase activity of ABCA4s showed two-fold stimulation by ATR, which correlated well with the previous studies with ABCA4 (9, 17, 43). In 1999, Sun *et al.* reported that different retinal isomers stimulated the ATPase activity approximately two-fold (9). Enzymatic data presented here showed that the soluble ABCA4s functions and its native membrane form in these *in vitro* assays.

To our knowledge, this is the first study that re-engineered such a large transporter protein with twelve TMDs to its soluble form and maintained the protein’s functional activities. Of note, refolding was not required to induce solubility or maintain enzymatic activity. The QNTY methodology, presented here, helped design the soluble form of a very complex membrane protein, ABCA4, with true solubility and enzymatic activities. Therefore, this method is likely applicable in creating soluble analogs of other complex transmembrane proteins.

## Materials and Methods

### Buffers and solutions

Buffer A contained 25 mM HEPES (pH 7.4), 130 mM NaCl, 1.5 mM MgCl_2_, 5% gycerol, 0.1% NP-40, and 1 mM DTT. Buffer B contained 0.1 mM glycine-HCl, (pH 3.0), Halt protease inhibitors (Thermofisher) in buffer A. Buffer C comprises 25 mM glycylglycine (pH 7.8), 15 mM potassium phosphate (pH 7.8), 15 mM MgSO_4_, 1 mM DTT. Buffer D comprises (1 μg/ml) Luciferase (Promega), and (0.15 mg/ml) D-Luciferin Potassium Salt (Biosynth/Carbosynth, CA) in buffer C. Buffer E consisted of 25 mM Tris-HCl (pH 7.5, 10% Glycerol, 0.1 mg/ml BSA, and 5 mM DTT.

### DNA sequence analysis

**DNA** sequencing confirmation of the recombinant plasmid ABCA4s was done at Eurofins MWG Operon LLC (Huntsville, AL).

### Computational protein modeling and structure analysis

The *in-silico* structures of native and soluble ABCA4 proteins were modeled with AlphaFold2 deep-learning-based protein modeling software using AlphaFold2_advanced. ipynb notebook in Google Colab (39, 40). The native ABCA4n and soluble ABCA4s protein models were superimposed and structurally compared in the PyMOL2 software (The PyMOL Molecular Graphics System, Version 2.0 Schrödinger, LLC). The Root-mean-square deviation (RMSD) was calculated to measure the structural alignment and overall conformational similarity.

### Baculovirus expression

ABCA4s was synthesized by vectorbuilder.com. ABCA4s was expressed in insect cells following the MultiBac^TM^ protocol (Geneva-Biotech, Geneva, Switzerland). Insect cells were cultured as monolayers, not suspensions. For protein production, virus-infected cells were harvested 62 hours post-infection. SFM-900 III serum-free medium (without antibiotics) was used to maintain insect cell lines.

### Assessment of protein solubilization

Expressed recombinant ABCA4s in the baculovirus expression system were extracted using I-PER buffer from ThermoFisher Scientific (Waltham, MA). To confirm the solubility of the protein, we analyzed the protein fractions by SDS PAGE for Coomassie R-250 staining and Western blot analysis (Figure 3). For standard Western blot, we used rabbit polyclonal anti-ABCA4 antibody (Abcam, ab72955) and secondary antibody was goat-anti Rabbit HRP (Abcam, ab205718).

### Protein purification

ABCA4s soluble fractions were dialyzed in buffer A for two hours at 4°C. Dynabeads protein G (ThermoFisher Scientific, Waltham, MA) coupled ABCA4 antibody was used for immunoaffinity purification of ABCA4s. We eluted the antibody bound ABCA4s using buffer B. The eluted fractions were immediately neutralized using pre-determined amount of 1 M Tris base followed by the addition of Buffer A. Protein concentrations were determined. Coomassie staining and Western blot analysis were performed to ascertain homogeneity of the protein (Figure 3C). Biological activity was assessed using ATR-stimulated ATPase assay. The purified ABCA4s retained full biological activity for at least 72 hours at 4°C and >3 months in aliquots at -80°C. Expression levels of ABCA4s in insect cells were uniformly high. We routinely purified ∼0.1 mg homogeneous ABCA4s from ∼1g of cell pellet. A large-scale, high-yield expression system is being developed.

### ATPase Assay

The ATPase assay was performed with purified ABCA4s in a 10 μl volume. The standard reaction mix contained ABCA4s enzyme (as indicated), 0.5 mM ATP, 5 mM MgCl_2_ in 1X buffer E with or without 40 μM ATR. Reactions were incubated at 37°C for one hour in the dark. Next, we diluted the ATPase reaction mix (1:30) in buffer C. Transferred 10μl of the diluted mixture to a 96 well-plate (Greiner, Germany) and 100 μl of buffer D was added and incubated at room temperature for 15 min in the dark. Luminescence was measured using a Clariostar Plate reader (BMG Labtech, Cary, NC). GraphPad Prism 9 was used to analyze the data.

## Acknowledgments

Authors gratefully acknowledge the support of this work by the Delaware Biotechnology Institute with an ARC-CAT grant, a fellowship grant to AAA by the SACM Cultural Mission, a fellowship grant to SC by the Republic of Turkey Ministry of National Education, and a Graduate Scholar fellowship grant to JSJ by the University of Delaware Graduate College.

## Supporting Information for the Manuscript

**Fig. S1.**
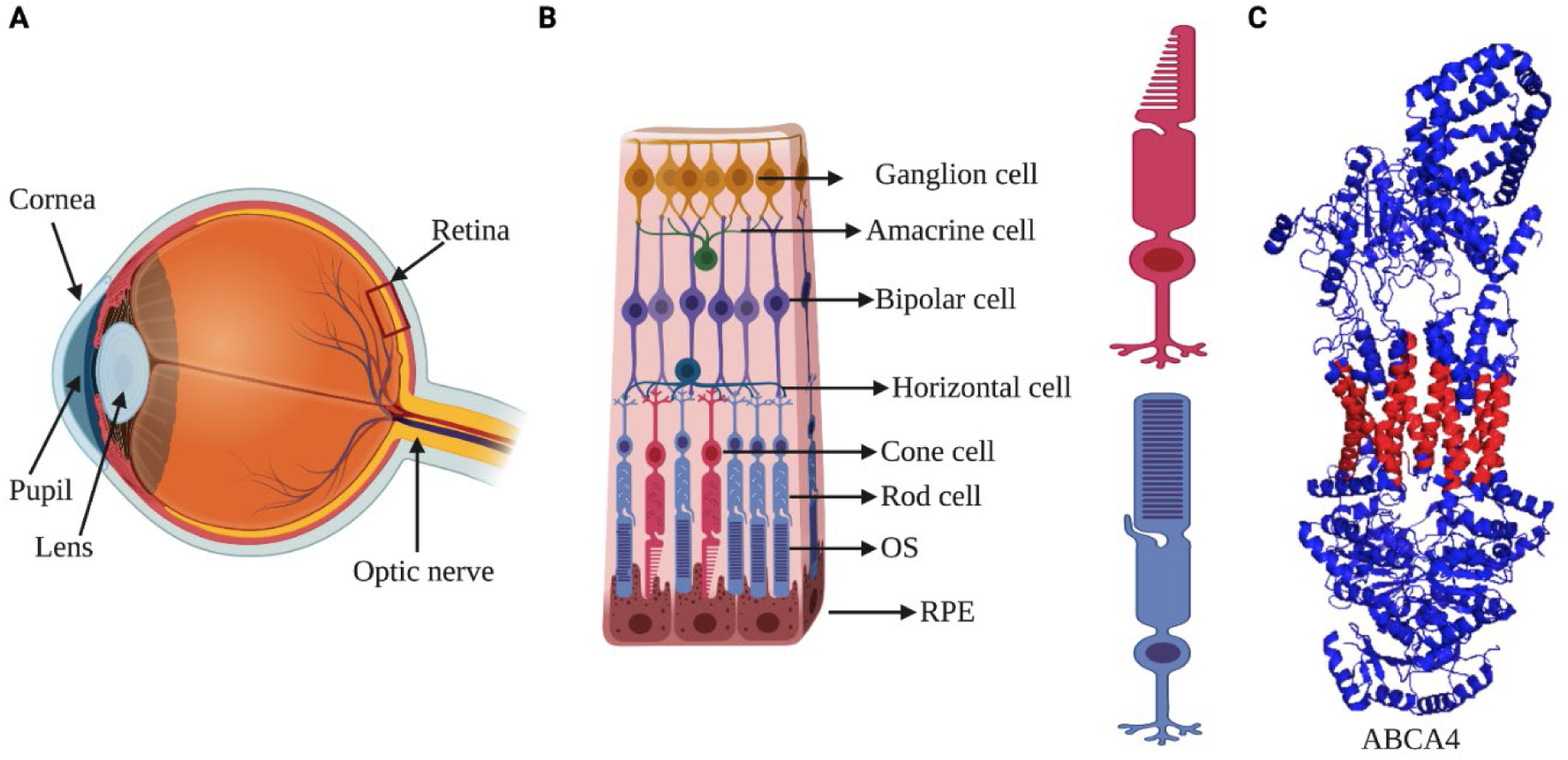
Location of ABCA4 in human retina. **A.** Cross-sectional diagram of the retina. **B.** ABCA4 is highly expressed in the rim region of the retina and is localized to the rod and cone outer segment discs. **C.** The structure of ABCA4 that solved by Liu F. *et al.* (2021), RCSB PDB: 7LKZ. Depicted on PyMol. The red color shows the 12 transmembrane segments (1). Created with BioRender.

**Fig. S2.**
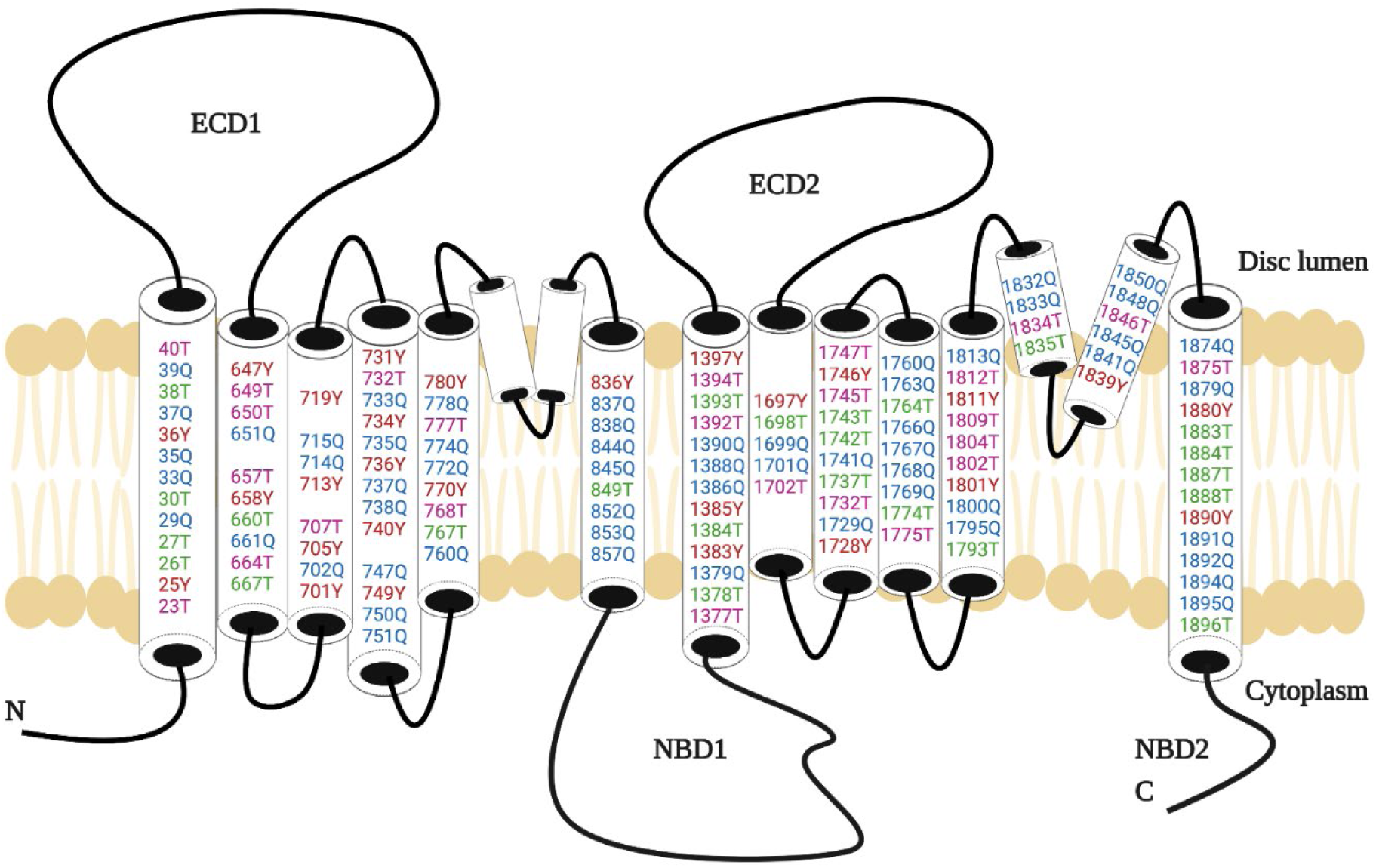
The topological structure of the 1st construct. QTY modification to the twelve transmembrane α helices, EH3 and EH4. QTY substitutions in the 12 transmembrane helices were as follows L→Q is shown in blue, I→T is presented in pink, V→T is colored in green, and F→Y is shown in red. Numbers indicate the sequence location where QTY substitution occurred.

**Table S1.**
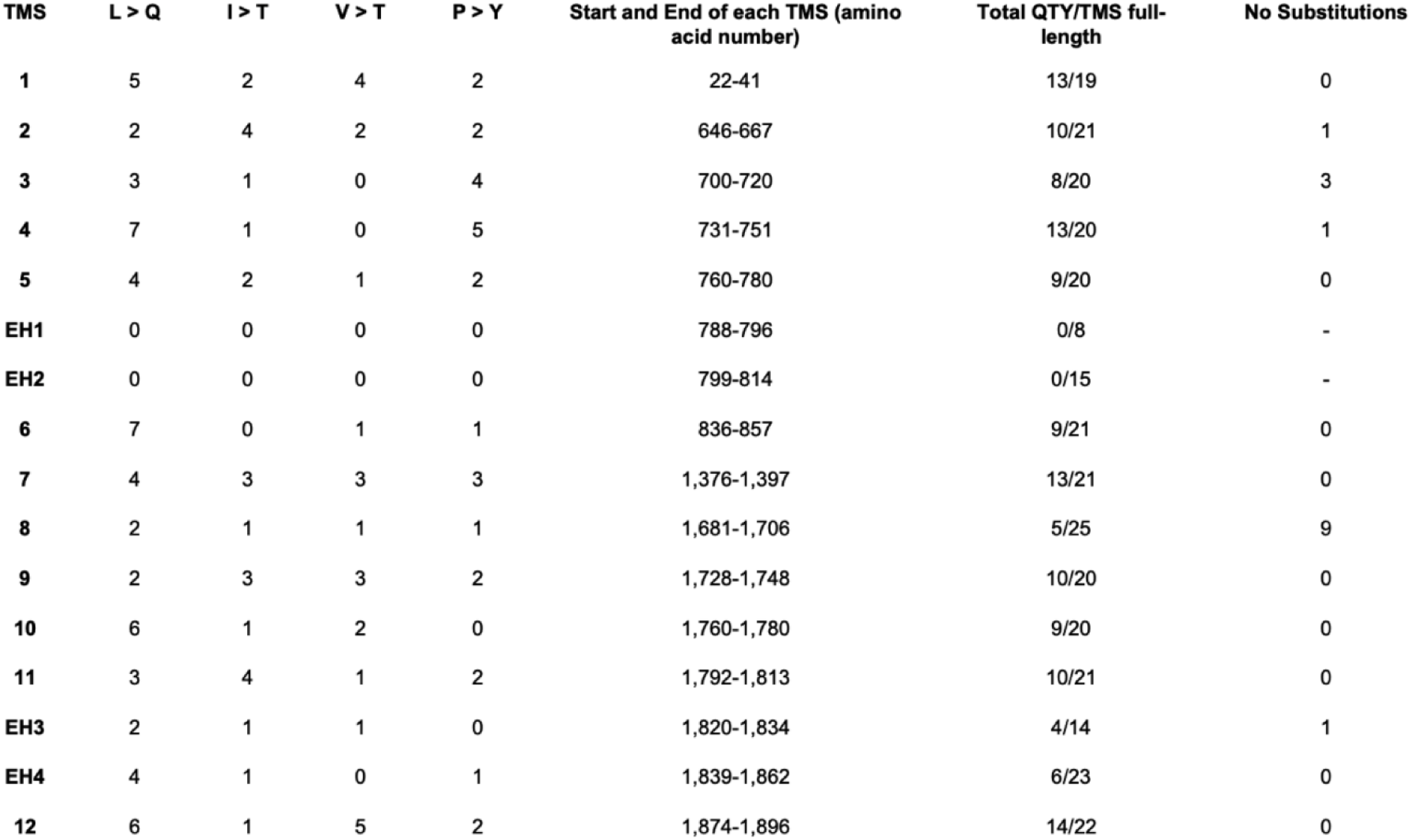
Summary of QTY modification to the 1st construct.

**Table S2.** Use of PolyPhen-2 Server To Select Between L → Q and L → N in 8 Leu Residues Substitution.

| TMDs | Amino Acid Substitution | PolyPhen-2 | Yes/No |
| --- | --- | --- | --- |
| 4 | L 747 Q | Benign | ✓ |
|  | L 747 N | Possibly Damaging | X |
|  | L 751 Q | Possibly Damaging | X |
|  | L 751 N | Benign | ✓ |
| 5 | L 772 Q | Possibly Damaging | X |
|  | L 772 N | Benign | ✓ |
|  | L 774 Q | Benign | ✓ |
|  | L 774 N | Probably Damaging | X |
| 6 | L 837 Q | Benign | ✓ |
|  | L 837 N | Possibly Damaging | X |
|  | L 845 Q | Possibly Damaging | X |
|  | L 845 N | Benign | ✓ |
|  | L 852 Q | Possibly Damaging | X |
|  | L 852 N | Benign | ✓ |
| 7 | L 1388 Q | Benign | ✓ |
|  | L 1388 N | Possibly Damaging | X |

**Fig. S3.**
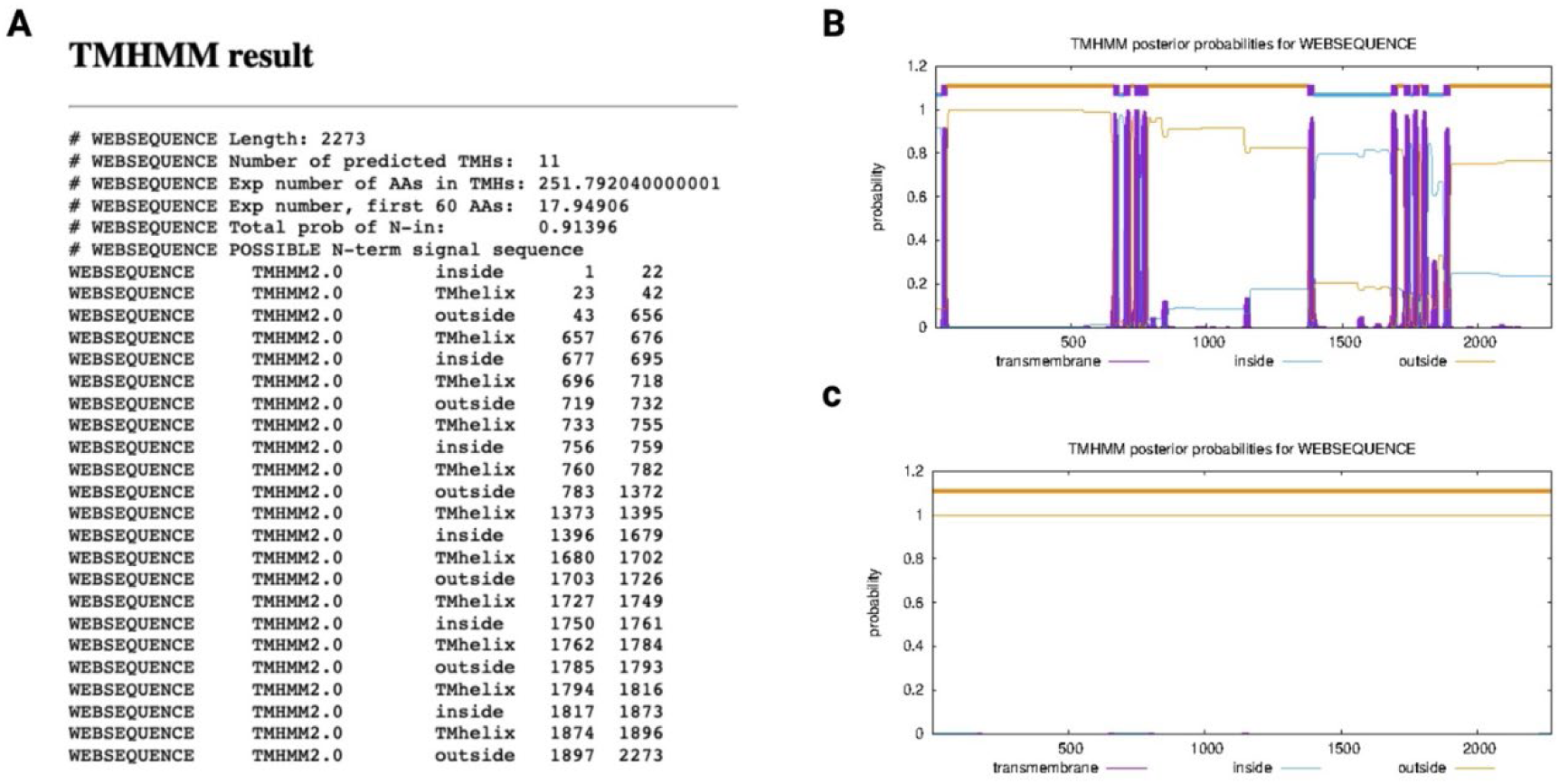
TMHMM 2.0 assessment of transmembrane α helices. **A**. Data of the native ABCA4 protein without QNTY modification, including the range of the intracellular segments, extracellular segments, and all the TM helix in the full-length ABCA4. **B.** Plot for the native ABCA4 shows the twelve transmembrane α-helices in pink, intracellular segments in blue, and extracellular segments in orange. **C.** After reengineering the full-length ABCA4 into its soluble form, ABCA4s, TMHMM software analysis was unable to detect any transmembrane helices, which is an indication of complete removal of most of the hydrophobic residues in the twelve transmembrane domain segments.

**Fig. S4.**
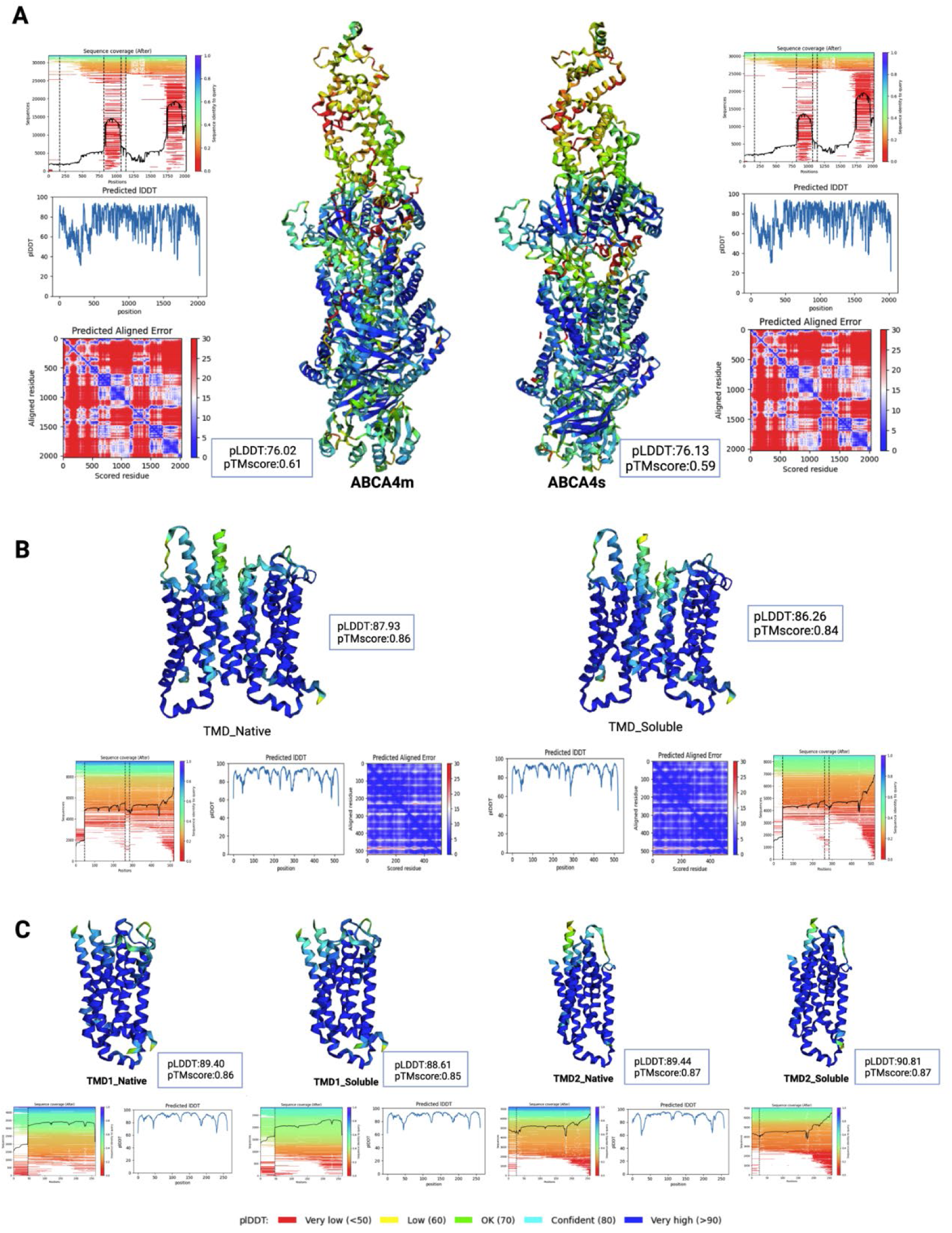
Quality scores of the AlphaFold2 protein models. The protein models with the sequence coverage, predicted aligned error, pLDDT, and pTM scores as produced in the AlphaFold2_advanced.ipynb notebook in Google Colab (2, 3) are presented here. The models were colored based on the pLDDT scores. All models were generated in high confidence with pLDDT> 75.

